# The *Synechocystis* ORF *slr0201* product is involved in succinate dehydrogenase-mediated cyclic electron transfer around PSI

**DOI:** 10.1101/2021.09.27.461999

**Authors:** Fusheng Xiong, Yang Yang, Xiyuan Fu

## Abstract

The *Synechocystis* sp. PCC 6803 open reading frame (ORF) *slr0201* was originally annotated as heterodisulfide reductase B subunit (HdrB). The *slr0201* encodes a 301-amino acid hypothetical protein with the predicted amino acid sequence significantly homologous to not only the HdrB from methanogenic bacteria, but also some novel succinate dehydrogenase C subunit (SdhC) found in Archaea and Campylobacter. Genetic manipulation via knocking-out approach created a *Δslr0201* mutant showing a *ΔsdhB*-like phenotype that was characterized by impaired succinate-dependent DCPIP reduction activities, reduced SDH-mediated respiratory electron transports, lower cellular contents of succinate and fumarate, slower KCN-induced increases in Chl fluorescence yield in the dark, and weak state 2/strong state 1 transitions, being indicative of a more oxidized PQ pool. In addition, slower re-reductions of the photosystem (PS) I reaction center P700 upon light-off were also monitored in the *Δslr0201*, indicating functional involvements of Slr0201 in cyclic electron transfer around PSI. Both photoautptrophical and photomixotrophical growth rates of the *Δslr0201* strain resembled to those of the wild type, but substantial growth deteriorations occurred when arginine (∼25 mM) or other two urea-cycle relevant amino acids (citrulline and ornithine) were added, which were attributed to generations and accumulations of certain hazardous metabolites. Based on the *Δsdh*B-resembling phenotype, in conjunction with its high sequence similarities to some archaeal SdhC, we proposed that the *slr0201* encodes a SDH function-relevant protein and is most likely the SdhC, a membrane anchoring subunit, which, while being genetically distinct from those in traditional bacterial SDH, belongs to the C subunit of novel archaeal SDH.

## Introduction

Bioenergetic electron transport in cyanobacteria is characterized by occurrences within a single cellular compartment of both photosynthetic and respiratory electron transports (PSET and RET). Unique structural continuity facilitates functional integrations of PSET and RET on cyanobacterial thylakoids, making the redox poising of plastoquinone (PQ) pool the most probable redox sensor on thylakoid membranes, being mutually regulated by not only PSET through photosystem (PS) II and I, but also stromal electron sources *via* RET flux into and out of the PQ pool. Models and evidence were established for interwoven PSET and RET in cyanobacteria and other photosynthetic bacteria (see reviews by *e.g.* Myers, 1986; Scherer, 1990; Vermaas, 2001; Peschek et al., 2004; Liu et al., 2012) which indicate that at least some components of the respiratory electron chain, such as NAD(P)H dehydrogenase, succinate dehydrogenase (SDH) and the cytochrome *aa_3_*-type terminal oxidase are localized on both cytoplasmic membranes and thylakoid membranes. In the thylakoid membranes, for example, some common electron carriers shared by PSET and RET processes include cytochrome *b_6_f* complex, PQ pool and soluble redox-active proteins (Vermaas, 2001; Durán et al., 2004; Peschek et al., 2004; Liu et al., 2012).

In cyanobacteria, it was evident that reducing equivalents for redox poising of the PQ pool can be derived through non-photochemical reactions. Three enzymes, succinate dehydrogenase (SDH), a NADPH-preferring type I (NDH-1) and a NADH-oxidizing type II dehydrogenase (NDH-2) are presumably main donors of respiratory electron flux into the PQ pool. In the cases of NDH-1 and NDH-2, sequence alignments through the genome of *Synechocystis* sp. PCC 6803 (hereafter *Synechocystis* 6803) predict the presences of *ndh*-1/*ndh*-2 gene-like open reading frames (ORFs) (Kaneko et al., 1996; Howitt et al., 1999). Genetic manipulations via deletions of the *ndh*-1gene-like ORF (*sll0223*) or *ndh*-2 ORFs (*slr0851*, *slr1743*, *sll1484*) created the Δ *ndh*-1 and Δ *h*-2 mutants. Both show a more oxidized PQ pool in darkness compared to the wild type, which is indicative of functional participations of these two enzymes in electron flux into the PQ pool (Howitt et al., 1999; Cooley et al., 2000; Ohkawa et al., 2000; Bernát et al., 2011). Turning to SDH, a traditional bacterial (*e.g. E. coli*) enzyme complex consists of a catalytic reaction center that includes a flavoprotein (SdhA) and an iron-sulfur protein (SdhB), both of which are highly conserved over a wide variety of organisms (Lancaster and Kroger, 2000; Lancaster, 2001). Additionally included are one or two membrane-bounded anchoring protein subunits (SdhC and /or SdhD or SdhE), being highly variable with respect to ligand and cluster components, which is, in fact, a classifiable basis over four or five groups of SDH (Lancaster and Kroger, 2000; Lancaster, 2001; Schäfer et al., 2002). In the genome of *Synechocystis* 6803, two *sdhB*-like ORFs (*sll1625* and *sll0823*) and one *sdhA*-like ORF (*slr1233*) are identified. Deletion of two *sdhB*-like ORFs generate a Δ*sll1625/*Δ*sll0823* (ΔSdhB_1_B_2_) double mutants with an even more oxidized PQ pool in the dark than in the Δ *ndh*-1 and Δ *ndh*-2 mutants (Cooley and Vermaas, 2001). This implies that in *Synechocystis* 6803, SDH not only functionally participates in redox poising of the PQ pool but also possesses relatively larger capacities for electron donation. While these results strongly indicate functional presence of SDH complex in *Synechocystis* 6803 and the important role of SDH in redox poising of PQ pool, sequence homologues to the membrane anchoring subunit(s) (SdhC and/or SdhD/SdhE) of a typical bacterial SDH are missing in genomics of *Synechocystis* 6803 according to the CyanoBase, therefore raising the question: how can the SDH complex function with only two catalytic subunits in absence of an essential membrane anchoring subunit?

Intriguingly, a hypothetical protein encoded by the ORF *slr0201* shows high sequence similarity to the C subunit of the novel archaeal SDH (Schäfer et al., 2002). This putative Slr0201 is originally annotated as heterodisulfide reductase B subunit (HdrB) [EC: 1.8.98.1] solely due to its sequence similarity to HdrB. Considering that (i) archaeal and bacterial HDR all consist of three subunits HdrA/B/C and their functional presence in *Synechocystis* 6803 has yet to be clarified and (ii) except for the *slr0201*, a putative *hdr*B given that sequence-based automatically-computational annotation were correct, neither *hdr*A-nor *hdr*C-gene like ORFs have been identified in the genome of *Synechocystis* 6803. Thus, similar questions raised as if this multi-subunit HDR complex does function in the *Synechocystis* 6803, how can it be only one relevant gene (*slr0201*) identified?

To address these uncertainties, genetic manipulations of the ORF *slr0201* were implemented, generating a *slr0201*knockout mutant (the *Δslr0201*). Physiological and biochemical analyses of SDH-relevant functions in the mutant were conducted and compared to the wild type and the *ΔsdhB_1_B_2_* strain that was previously constructed (Cooley et al., 2000; Cooley and Vermaas, 2001). The *Δslr0201* strain illustrated a *ΔsdhB_1_B_2_*-like phenotype in terms of impaired succinate dehydrogenase activity, reduced substrate (succinate and fumarate) levels, low SDH-mediated respiration rate, more oxidized PQ pool and slower re-reduction of P700 in the dark. The *ΔsdhB_1_B_2_*-resembling phenotype of the *Δslr0201* strain in conjunction with the sequence presence of the *sdh*A/B-like genes and absence of the *hdr*A/C-like genes in the genome of *Synechocystis* 6803, high sequence similarities with the C subunit of the “non-classical” archaeal SDH, and the *in vivo* overexpression of the thylakoid-associated Slr0201 (Xiong and Vermaas, unpublished) suggested that the *slr0201* encodes a SDH function-relevant protein, rather than the HdrB. Most likely, the Slr0201 acts as a membrane anchoring subunit of SDH complex. While this putative protein is distinct from the SdhC from traditional bacterial SDH, it highly resembles the novel SdhC from “non-classical” group of archaeal SDH.

## Materials and methods

### Construction of slr0201 deletion mutants and growth conditions

To create a *slr0201* knockout mutant, two *slr0201*-flanking regions (up- and down-stream) of the *Synechocystis* 6803 genome, according to CyanoBase (http://www.kazusa.or.jp/cyano), were amplified by PCR using two pairs of forward and reverse primers that introduced unique restriction sites, EcoRI/BamHI and BamHI/PstI, respectively (Fig.1A). The up- and down-stream fragments were then ligated with a 1.3 kb kanamycin resistant cassettes at the engineered BamHI sites and cloned into the pUC19 using the EcoRI and PstI restriction sites. The generated plasmid named Δ*slr0201*-Km was used for transforming the wild-type *Synechocystis* sp. strain PCC 6803 which was detained previously (Cooley et al., 2000). The transformants were segregated under an increased kanamycin selection pressure. Segregation analysis was performed by PCR using primers specific for the sequences of the flanking regions of the *slr0201* being deleted. The exclusive presence of a band corresponding to the inactivated gene was taken as evidence of full segregation (Fig. 1B). In addition, PCR was also performed by using primer-recognizing sequences inside the wild-type sequence that was deleted in the mutant (data not shown). Double mutants were obtained by transforming the PSI-deficient strains or the *Δslr1022* with the plasmid pΔ*slr0201*-Km.

**Figure 1.**
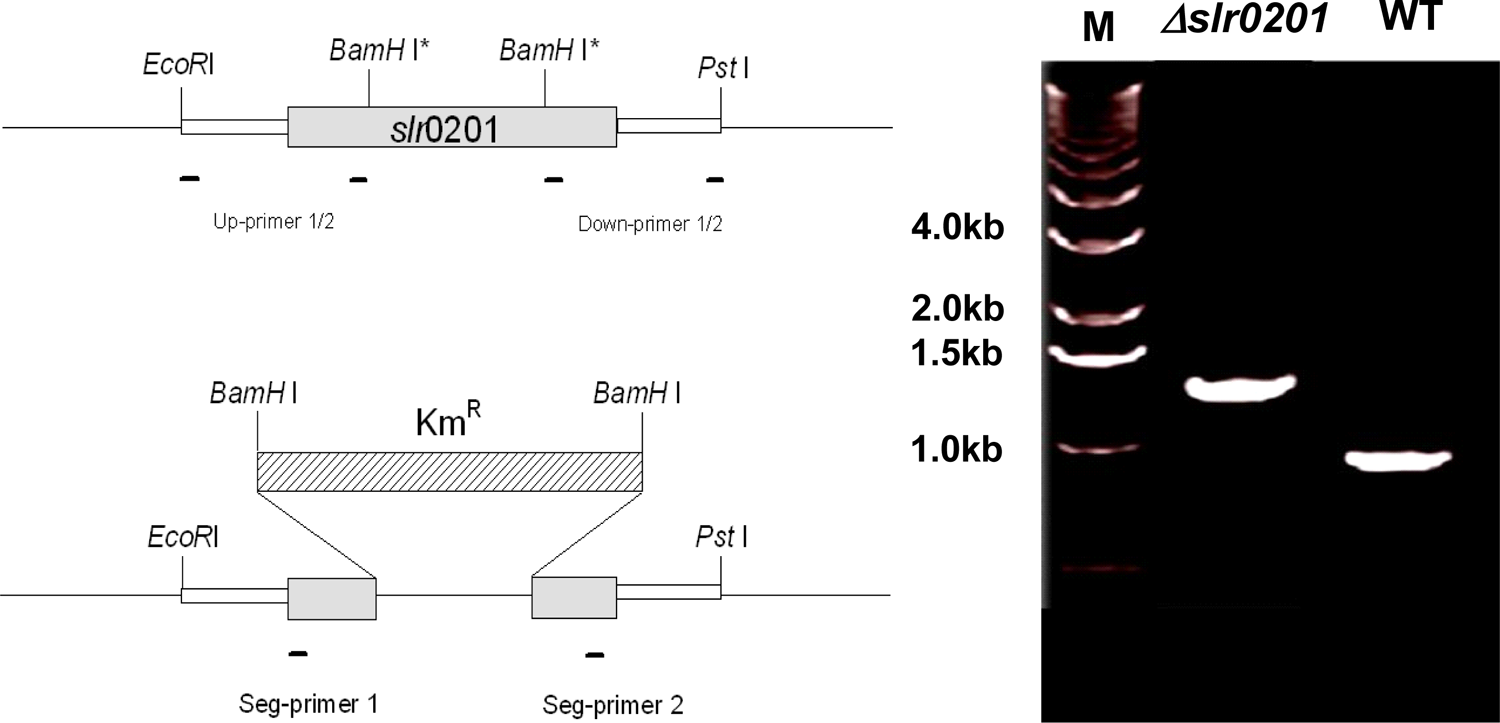
(**A**). Schematic illustration of *Synechocystis* sp. PCC 6803 genomic DNA encompassing the open reading frame Δ*slr0201* (thick gray square). The sequence between the two *Bam*H I sites in the original ORF *slr0201* was replaced by the Km resistance cassette (1.3 kb) from pUC4K. (**B**). Agarose gel electrophoresis profiling of PCR products amplified using genomic DNA extracted from the WT strain (WT) and the Δ*slr0201-deletion* mutant (*Δslr0201*) indicates a complete segregation of the mutant. Lane M. 1 kb DNA ladder.

The wild type and mutant strains of *Synechocystis* 6803 were grown at 30 °C in liquid BG-11 medium buffered with 10 mM TES-NaOH (pH 8.0) and illuminated at a light intensity of 50 µmol photons m^-2^ s^-1^. For photomixotrophic growth the medium was supplemented with 5 mM glucose. For growth on plates, 1.5% (w/v) agar and 0.3% (w/v) sodium thiosulfate were added to BG-11 medium with appropriate antibiotics to which a particular strain was resistant due to the presence of antibiotic resistance cassette introduced with gene inactivation. Antibiotic concentrations used were 25 µg mL^-1^ of kanamycin, 10 µg mL^-1^ of erythromycin, and 25 µg mL^-1^ of spectinomycin. When appropriate, arginine and other amino acids (citrulline and ornithine) were added to the growth medium as specified in the text (see the result section).

### Succinate-dependent DCPIP reduction in the isolated membranes

Thylakoid membranes were prepared from the wild type and the Δ*slr0201* strains growing in the mid-exponential phase as described previously (Yu and Vermaas, 1993). Succinate:quinone reductase activity was determined by monitoring its capacity to catalyze succinate-dependent reduction of quinone-stimulated dichlorophenolindophenol (DCPIP) following Yang et al. (1998). The assay was performed at 37 °C with a Hewlett Packard HP8452 diode array spectrophotometer. Membranes (50 μL, ∼ 25 μg Chl) were added to 1 mL assay mixture containing 0.1 M sodium/potassium phosphate (pH 7.5), 50 µM DCPIP, 20 mM succinate, 1 mM EDTA, 0.01% Triton X-100, 10 μM KCN, and 25 mM of 2,3-dimethoxy-5-methyl-6-geranyl-1,4-benzoquinone (Q_2_). The reduction of DCPIP was monitored by measuring the absorbance decrease at 598 nm. The millimolar extinction coefficient used was 21 mM^-1^ cm^-1^.

### Measurements of respiratory O_2_ consumption

Rates of respiratory O_2_ consumption were measured with whole *Synechocystis* 6803 cells at 30 °C using a Clark-type electrode (Hansatech, UK) as previously described (Howitt and Vermaas, 1998). In brief, fresh cyanobacterial culture (OD_730_ ≈ 0.6) was spun down and re-suspended in 25 mM Hepes/NaOH, pH 7.0 at ∼ 10 μg Chl mL^-1^. After a 3-5 min thermal equilibration, respiratory O_2_ uptakes in darkness were initiated by adding 5 mM glucose (the final concentration) and were recorded over a 15 min time frame.

### Determinations of succinate and fumarate content

Quantitative determination of organic acid content in *Synechocystis* 6803 cells followed the procedure described previously (Cooley and Vermaas, 2001) with minor modifications. To extract organic acids from cyanobacterial cells, 1 L of fresh culture with OD_730_ = 0.6-0.8 was harvested by centrifugation and sucked onto a filter (0.45 μm pore size). Cells were quickly rinsed three times with ice-cold ddH_2_O within 30-60 s. The filter with the cell pellet was then immersed in 5 mL of 5% (w/v) perchloric acid for 10 min. After removing the filter, 2 M potassium bicarbonate was added drop-wise to the extract to bring pH to 3.2-3.5. The supernatant was finally collected after the extract was centrifuged for 30 min at 48,000 x g. Organic acids in the cell extract were purified using Sep-Pak cartridge (Waters Associates, Deerfield, MI) which was preconditioned following the manufacturer’s instruction. Gas chromatography-mass spectral (GC/MS) analysis was performed following the protocol (Cooley et al., 2000) using a Shimadzu 17-A gas chromatograph and a Shimadzu QP5000 mass spectrometer linked to a data processor (Class-5K GC/MS software; Shimadzu).

### Chlorophyll fluorescence and whole-cell state transition analysis

The level of chlorophyll (Chl) fluorescence induced by KCN in darkness and Chl fluorescence during whole-cell state transitions were monitored at room temperature using a modulated PAM fluorometer (Walz, Effeltrich, Germany). Cells were harvested at mid-exponential growth phase (OD_730_ ≈ 0.6) with low-speed centrifugation at 2800 x g for 5 min. The pellet was then re-suspended in 10 mM Hepes/NaOH (pH 7.0) at a Chl concentration of 5 µg mL^-1^. To monitor KCN-induced Chl fluorescence increase in darkness, cell suspension was first dark-adapted for 5 min prior to a 15-s measurement of the initial fluorescent level (Fo). Immediately afterwards, the measuring light was turned off and 1 mM KCN (final concentration) was added. The fluorescence yield was monitored immediately by turning on the measuring light again at 15-s intervals (10 s off, 5 s on) for the initial 60 s, and then at 35 s interval (30 s off, 5 s on) for an additional period of 10 min. The measuring light was set at its minimum (< 0.01 µmol photons m^-2^s^-1^) in order to avoid any noticeable actinic effect. The whole-cell state transitions were monitored followed Wilson et al. (2006) with illumination of blue light (450 nm, Corning 4.960) at 50 µmol photons m^-2^ s^-1^, orange light (601 nm, filter Balzer B-40 599 10) at 45 µmol photons m^-2^ s^-1^, and white light at 150 µmol photons m^-2^ s^-1^ used as actinic light for steady-state photosynthesis. Maximum Chl fluorescence level (Fm or Fm’) was determined using saturating light pulse (3500 µmol photons m^-2^ s^-1^) produced by a KL-1500 lamp (Schott, Germany).

### Measurements of P700 oxidation and re-reduction in vivo

Kinetics of P700 oxidation and dark rereduction were measured at room temperature with the concentrated whole-cell suspension (see below) by monitoring both initial slope and half-time of absorbance changes at 820 nm using a modulated PAM PL102 fluorometer (Walz, Effeltrich, Germany) as described elsewhere (Schreiber et al., 1988; Herbert et al., 1995). Transient changes of absorbance at 820 nm upon flashing- on/off were taken as proportional to changes in the redox status of P700. Specifically, the flash-on induced initial oxidation rate is thought to be proportional to the efficiency of excitation energy delivered from PSI light-harvesting antenna to P700, while transient decay (P700^+^ rereduction) rate after flash-off is proportional to the rate of electron flows into P700 via various electron transfer pathways.

To prepare cell suspension for P700 oxidation-reduction measurement, fresh *Synechocystis* cultures were harvested at exponential growth phase (OD_730_ ≈ 0.6) by low-speed centrifugation (2800 x g, 5 min). After washing with 50 mM Hepes/NaOH buffer (pH 7.0), the cell pellet was re-suspended and concentrated to OD_730_ ≈ 40. Before measurement, a 30-s initial illumination with far-red light (FR) was applied to reduce ferredoxin and the cell suspension was then kept in darkness for 10 min, during which, a small volume (5-10 μL) of 3-(3,4-dichlorophenyl)-1,1-dimethylurea (DCMU) or 2,5-dibromo-3-methyl-6-isopropylbenzoquinone (DBMIB) was added. Actinic light (560 μmol m^-2^ s^-1^) was provided by a tungsten projector lamp with multiple-turnover time settings. Flashing illumination periods (10-300 ms) were controlled by a custom-made device equipped with a UNIBLITZ-26L2A0T5 electronic shutter triggered by electronic signal derived from the PAM PL102 units. Unless otherwise specified, five flash-induced transient signals at 820 nm were averaged for each measurement.

## Results

### Growth of the Δslr0201 strain is similar to the wild type but sensitive to arginine and other amino acids in urea cycle

Under normal light intensity (50 μmol photons m^-2^ s^-1^) at 30°C, the Δ*slr0201* strain had a doubling time of 12.5 ± 1.5 for photomixotrophic growth, and 19.0 ± 2.1 h for photoautotrophic growth, which are similar to the wild type (data not shown). No significant effects of succinate and fumarate supplements on growth of both the wild type and the *slr0201* strain were observed. However, growth of the Δ*slr0201* strain was sensitive to arginine supplemented in the BG-11 medium. Growth deterioration appeared after 2-3 weeks of growth in the presence of arginine (Fig. 2A) even the initial growth of the Δ*slr0201* cells was, in fact, unaffected by arginine supplements as illustrated by successes in initial establishments of colonies. Growth deterioration was more significant at higher concentration (50 mM). The Δ*slr1022* strain, an arginine auxotroph (Quintero et al., 2000) was included as a positive control. Note that, in the connecting region between the areas #2 and #4 (Fig. 2A), the more the Δ*slr1022/* Δ*slr0201*cells were close to the Δ*slr0201*, the more these cells suffered. One explanation is that during growth of the Δslr0201 with the arginine supplement, certain kind of hazardous subtract was generated and as growth continued, this hazardous subtract was accumulated and diffused across the plate. The Δ*slr1022/*Δ*slr0201* mutant exhibited moderate arginine sensitivities, which could be a compromise between the Δ*slr1022-mediated* arginine auxotroph and the Δ*slr0201*-associated arginine vulnerability. Two other amino acids in the urea cycle, citrulline and ornithine show even more significant deteriorative effects on the growth of SDH mutant strains (Fig. 2B). Interestingly, growth deteriorations caused by citrulline and ornithine in the Δ*slr0201* were totally prevented with additional arginine supplement at low concentration (2 mM). For the *sdhB_1_B_2_*, however, only very limited alleviations occurred. The arginine-sensitive phenotype was further refined in liquid cultures. While during the initial culture all four tested strains showed successful growths even with 25 mM arginine addition, growth deteriorations appeared in the two SDH mutants as subculture continued (Fig. 3A). Similar to the plate growth (Fig. 2), growth deteriorations caused by arginine are concentration dependent. For the Δ*slr0201* (Fig. 3B), for example, during the 2^nd^ subculture, significant growth deterioration appeared only at 25 mM, the deterioration could be clearly seen at 5 mM during the 3^rd^ subculture.

**Figure 2.**
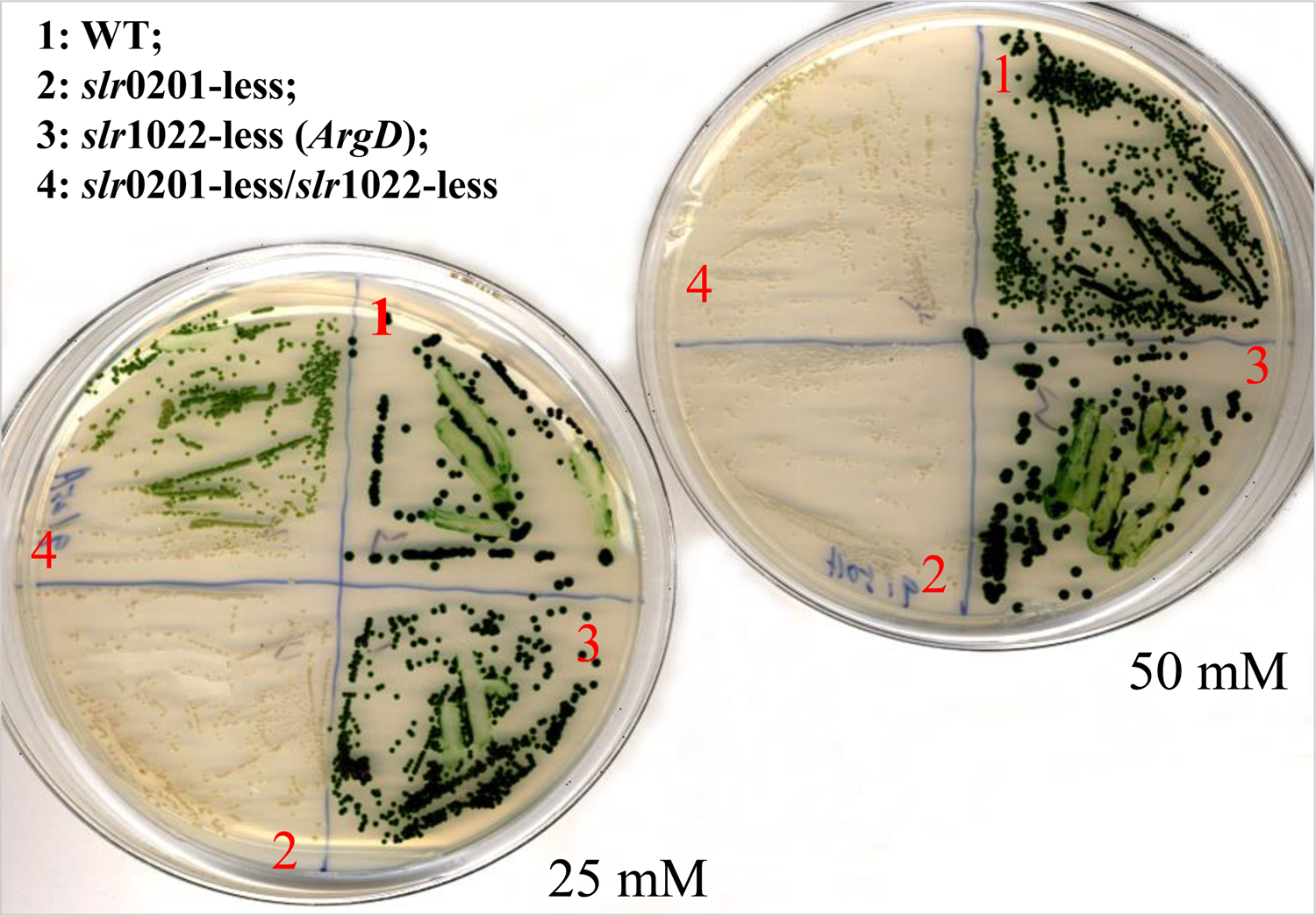
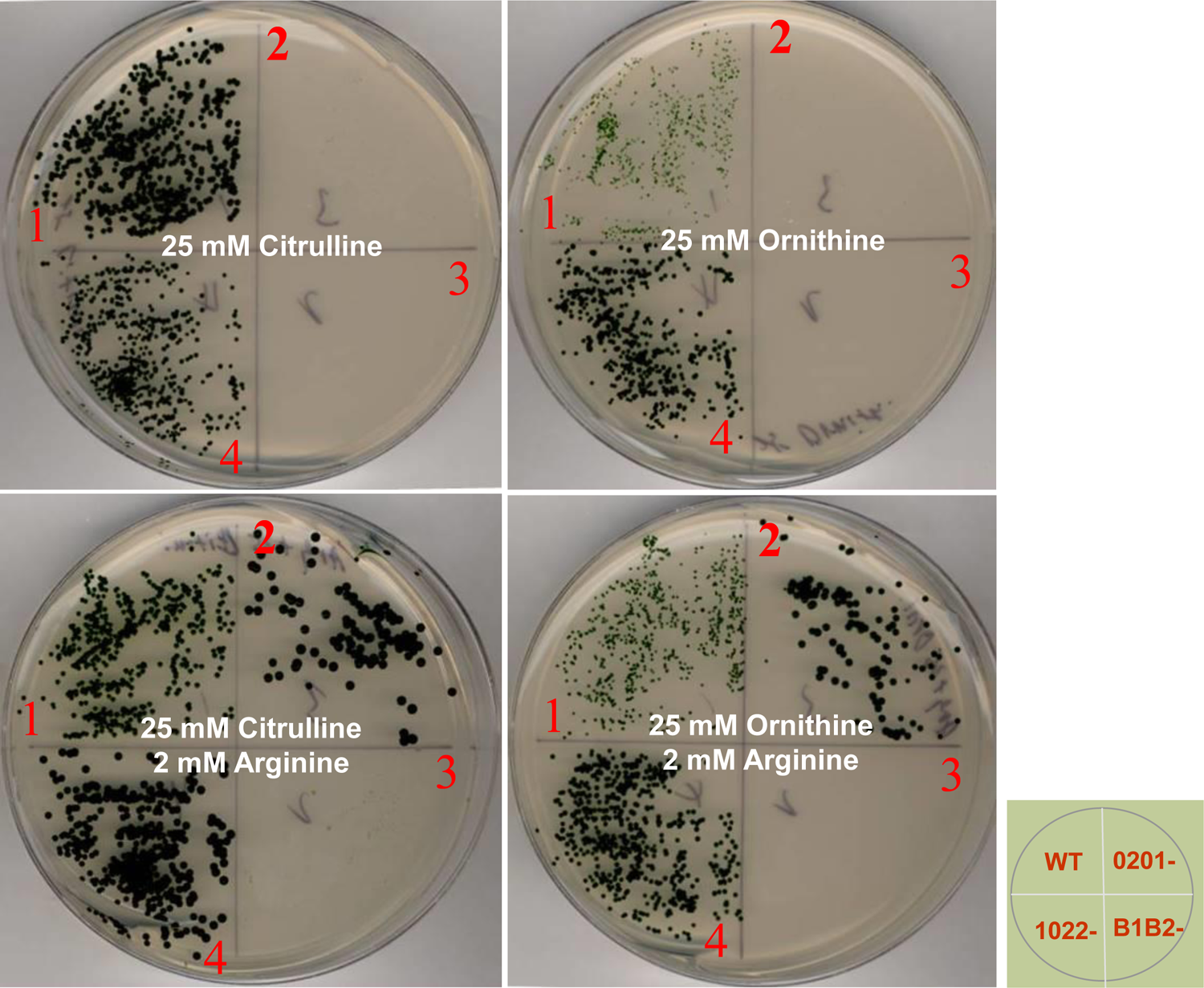
(**A**). Photoautotrophic plate growths of *Synechocystis* sp. PCC 6803 strains in the BG-11 plate medium supplemented with 25 and 50 mM arginine. (**B**). Photoautotrophic plate growths of *Synechocystis* sp. PCC 6803 strains in the BG-11 plate medium supplemented with 25 mM citrulline or 25 mM ornithine with or without 2 mM arginine addition. The photos were taken after 3-week growth.

**Figure 3.**
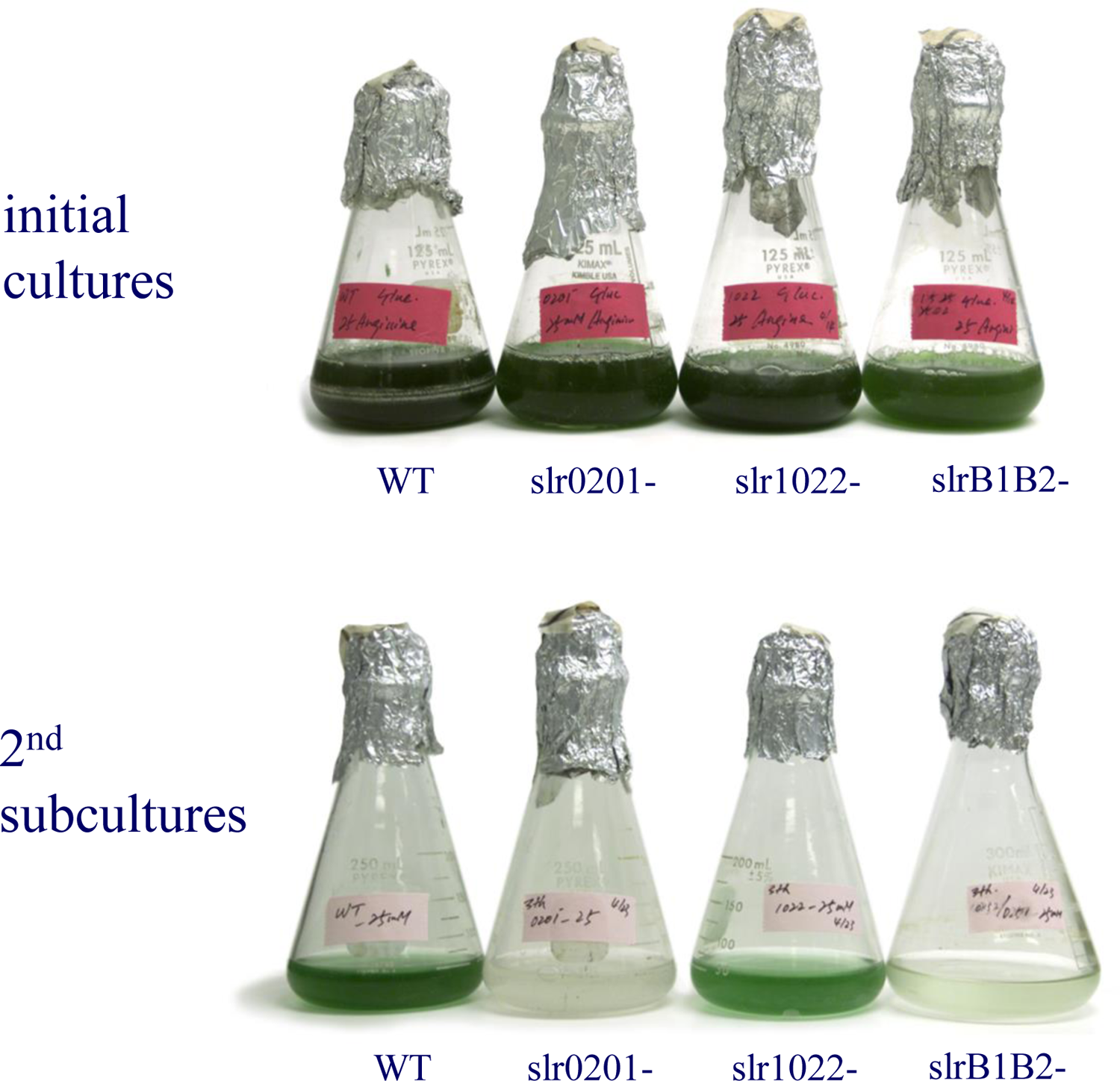
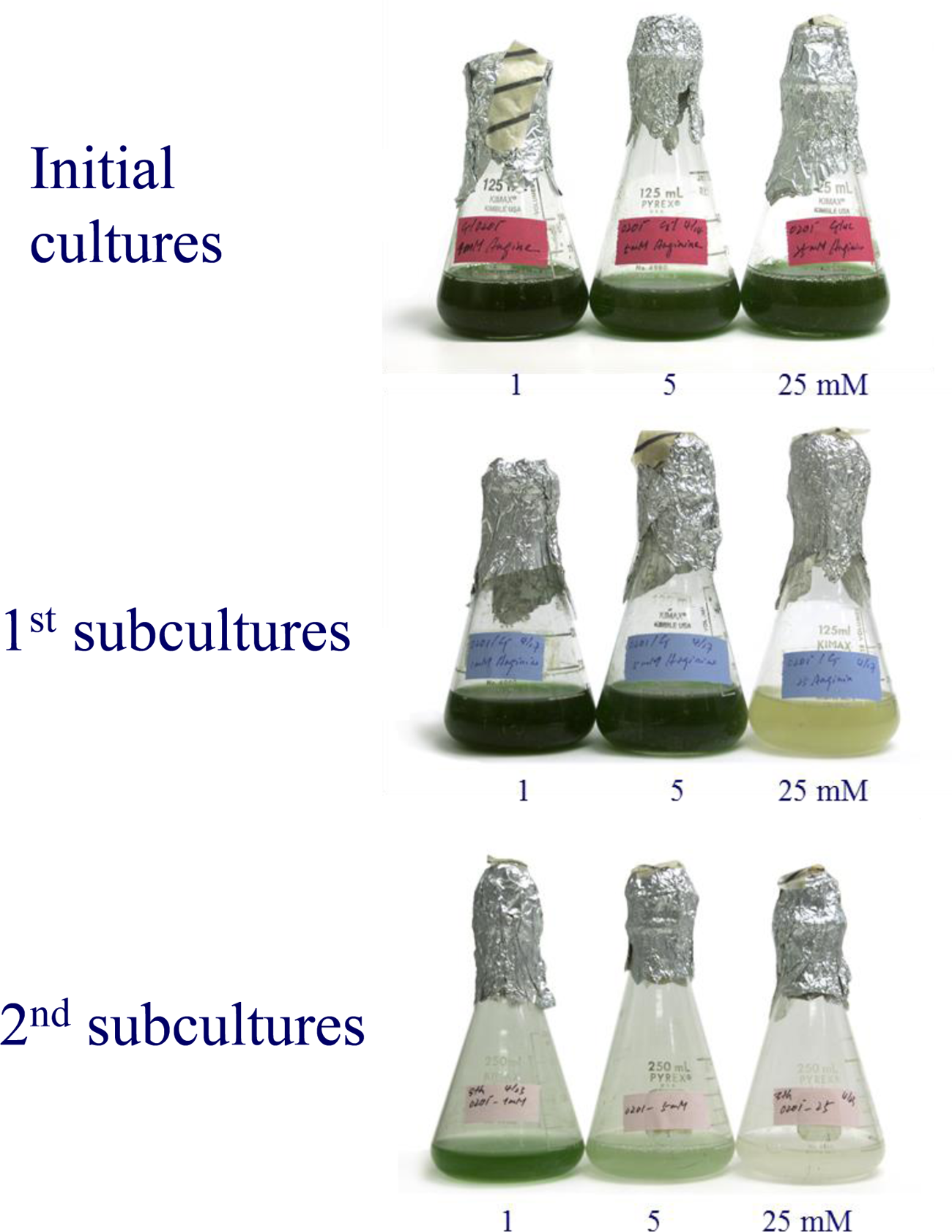
Arginine sensitivities of *Synechocystis* sp. PCC 6803. (**A**) The initial cultures of the wild type, the Δ*slr1022* and the Δmedium with addition of 25 mM arginine followed by three sequential 2-week subcultures. (**B**) Effects of 1, 5 and 25 mM arginine addition on the growths of the Δ*slr0201* strain in the BG-11 medium.

### Deletion of the ORF slr0201 results in significant losses of SDH activity in vitro

Measurement of succinate-dependent DCPIP reduction provides an estimate of the *in vitro* SDH activity (Yang et al., 1998). Membranes isolated from the Δ*slr0201* strain have a rate of succinate-dependent DCPIP reduction at 5.8 ± 0.7 μ being 52% lower compared to the wild type (Fig. 4). The reductions tended to be less pronounced when compared to the SDH activities in the Δ*sdhB_1_B_2_* strain. This result indicates that the Slr0201 is essential to a functioning SDH complex. However, the Slr0201-overexpressed *Synechocystis* cells did not show any significant enhancements of the SDH activity. An explanation could be that while the Slr0201 presumably acts as a membrane-anchoring subunit which is essential for SDH activity, extra Slr0201 copies alone might not be sufficient to result in any direct enhancements on SdhA/SdhB-catalyzing succinate reductions.

**Figure 4.**
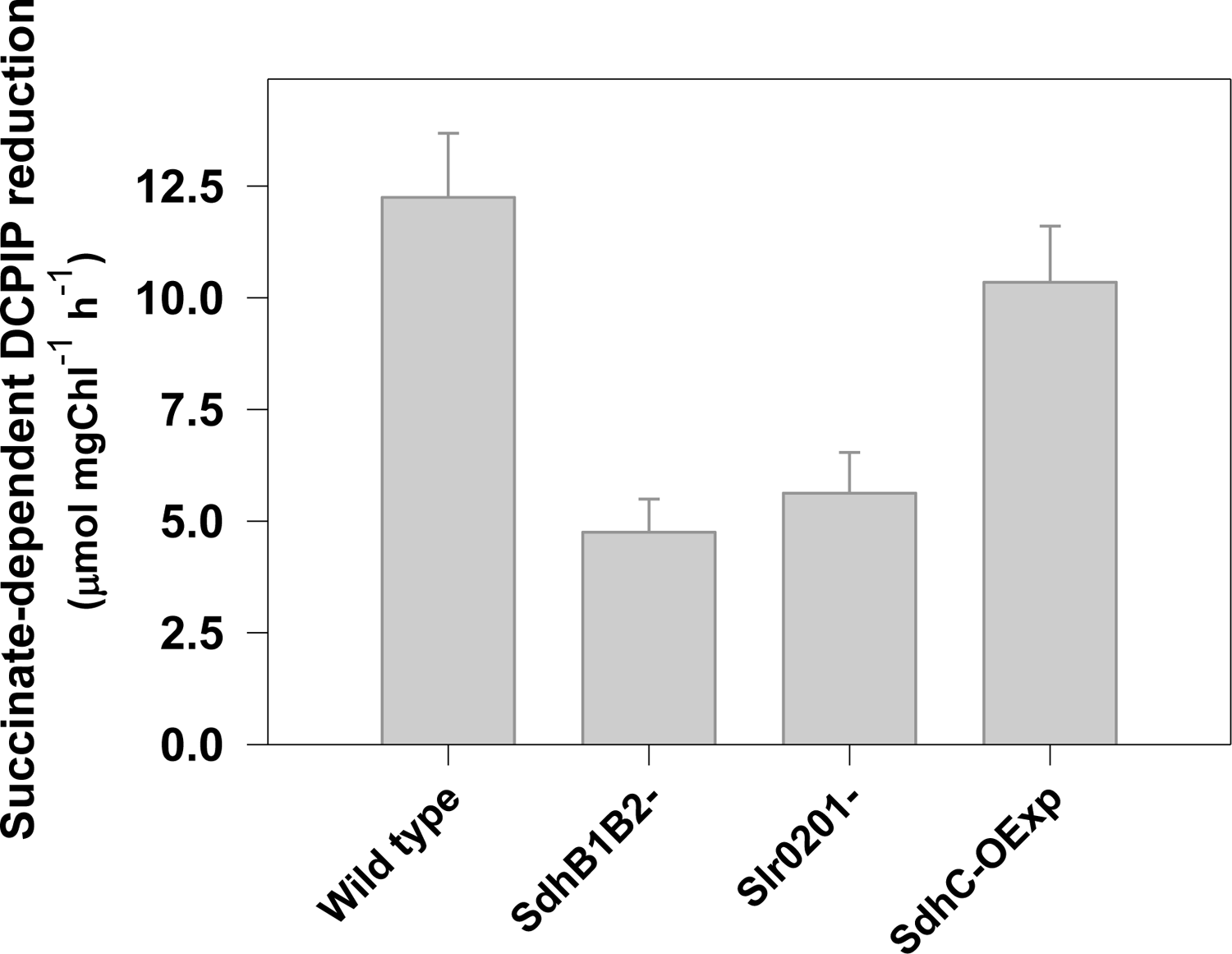
Succinate-DCPIP dependent SDH activities in *Synechocystis* sp. PCC 6803 strains. The values represent the averages and the error bars show ± SD (n=3). Both were calculated from three independent experiments. Each sample was adjusted to a Chl concentration of 5 µg Chl mL^-1^

### Fumarate and succinate levels are low in the Δslr020 strain

To determine succinate and fumarate content in the whole cells, methanol extraction of polar metabolites from whole cells was subjected to derivatization with TBDMS and followed by consequent quantification using GC/MS (Table 1). In the wild type cells, fumarate level was 0.09 ± 0.05 µmol g^-1^ FW. This level was ∼ 6-fold lower than the succinate level (0.54 ± 0.08 µmol g^-1^ FW). Deletion of the *slr0201* resulted in tremendous reductions in cellular contents of both succinate and fumarate. In the Δ*slr0201* mutant, fumarate level was 0.02 ± 0.01µmol g^-1^, which was almost 4.5-fold lower as compared to the wild type. Succinate level was also low in the mutant, being 0.29 ± 0.06 µmol g^-1^ although the reduction (∼ 46% lower than in the wild type) was not as pronounced as in fumarate

**Table 1.** Succinate and fumarate levels in *Synechocystis* sp. PCC 6803 strains

| Strains | Succinate | Fumarate |
| --- | --- | --- |
|  | (μmol g<sup>-1</sup> FW) |  |
| Wild type | 0.54±0.08 | 0.09 ± 0.05 |
| *Δslr0201* | 0.29±0.06 | 0.02 ± 0.01 |
| *ΔsdhB<sub>1</sub>B<sub>2</sub>* | 0.13±0.03 | ND <sup>a</sup> |

ND<sup>a</sup>: Not determined
The values represent the average ± SD calculated from three independent experiments.

### Respiratory O_2_ consumption is low in the photomixotrophically grown Δslr0201 strain

The overall respiratory electron transfer rate was estimated by measuring oxygen consumption of the cell suspension in darkness in the presence of 5 mM glucose. The results were presented as means ± SD (n=4). Respiratory rate was 44 ± 10 μmol O_2_ mg Chl^-1^ h^-1^ in the wild type and 18 ± 7 μmol O_2_ mg Chl^-1^ h^-1^ in the Δ*slr0201* strain. The latter was ∼ 60% lower than the former. The Δ*sdh*B_1_B_2_ strain consumed O_2_ at an even lower rate, being 11±4 μmol O_2_ mg Chl^-1^ h^-1^, which was only ∼ 25% of the respiratory rate in the wild type.

### Electron flows into the PQ pool through respiratory pathway are slow in the Δslr0201 strain

Monitoring Chl fluorescence yield provides insight into the redox state of Q_A_, a primary electron-accepting PQ in PSII, which is closely correlated with the redox state of the PQ pool. With addition of KCN to block electron flux out of the PQ pool via terminal oxidases, electron flows into the PQ pool could cause its reductions. If the PQ pool is sufficiently reduced, the Q_A_ will then become reduced due to an efficient redox equilibrium between the Q_A_ and the PQ pool (K_eq_= 23). Thus, the redox state of the PQ pool can be at least qualitatively determined by monitoring fluorescence yield following minimal PSII excitations. In the wild type (with the PSI-less background), upon addition of KCN in darkness, a rapid increase in fluorescence yield was evident (Fig. 5, the gray line), which reflects accumulations of the reduced Q_A_ due to over-reduction of the PQ pool. The PSI-less strain was used because (1) it had significantly higher variable fluorescence yield, and (2) excitations of PSI by measuring light will not result in significant oxidization of the PQ pool. Compared to the PSI-less strain set as a control, the *Δslr0201*/PSI-less and the Δ*sdhB_1_B_2_*/PSI-less strains both showed a substantial delay in initial increases of the KCN-induced fluorescence yield (Fig. 5, the dark gray and black lines). The immediate/transient rises in fluorescence yield upon addition of KCN were interpreted as a result of inhibition of oxidases by KCN such that electron flows out the PQ pool was blocked. After that, slower increases in variable fluorescence over a period of ∼ 3-5 min reflect a gradual reduction of the PQ pool mediated by partial reductions of the Q_A_. Note that a longer half time (t_1/2_) of 85 s to reach maximum fluorescence in the *slr0201* strain as compared to a t_1/2_ of 23 s in the wild type indicates that in the Δ*slr0201* strain, respiratory electron flows into the PQ pool were impaired. These results support the previous view that the SDH-mediated respiratory electron fluxes into the PQ pool greatly contribute to cyclic electron transfer around the PSI (Cooley and Vermaas, 2001).

**Figure 5.**
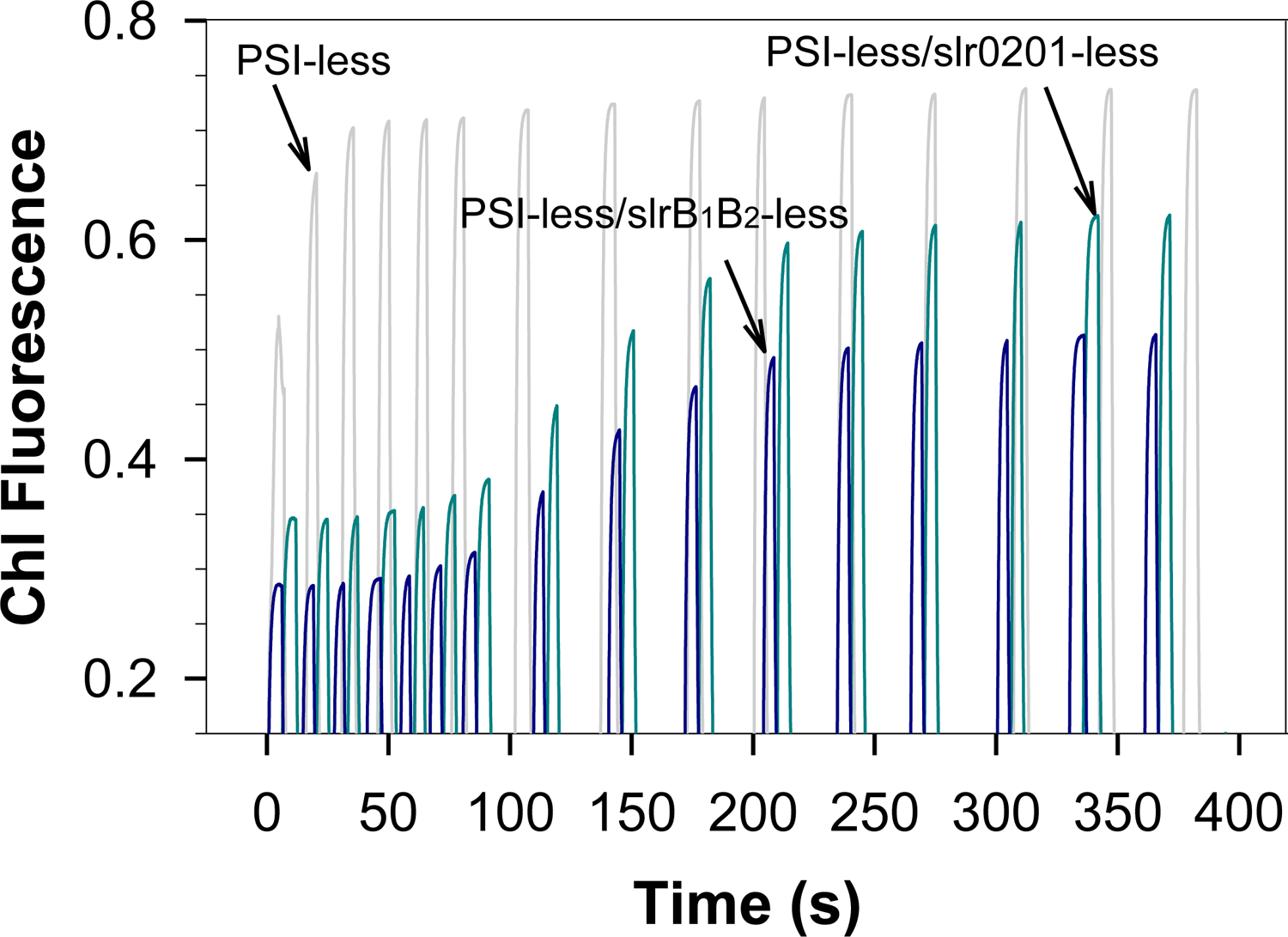
KCN-induced increases of fluorescence yield in darkness in *Synechocystis* sp. PCC 6803 strains. KCN was added at time zero and the measuring flashes were triggered in order to monitor fluorescence changes. See the M&M section for the details. For better observations of variable fluorescence yield, all strains have the PSI-less background.

### Chl Fluorescence quenching and light-/inhibitor-driven state transitions in the Δslr0201 strain

Potential involvements of the Slr0201 in the SDH-mediated electron fluxes through the PQ pool were further evaluated by monitoring Chl fluorescence quenching and light-/inhibitor-driven state transition. During steady-state photosynthesis Chl fluorescence kinetics in the *Δslr0201* strain was characterized by markedly higher steady-state fluorescence yields (Fs) along with substantial low steady-state variable fluorescence (Fv’) although no significant alternations occurred in neither Fm nor Fo (Fig. 6). Similar observations were previously reported in the PSI-light acclimated plants (Pfannschmidt et al., 2001), the M55 mutant (Mi et al., 1992) and the VIPP-less mutant (Kroll et al., 2001), which was ascribed to impaired electron transfer capacities in these plants.

**Figure 6.**
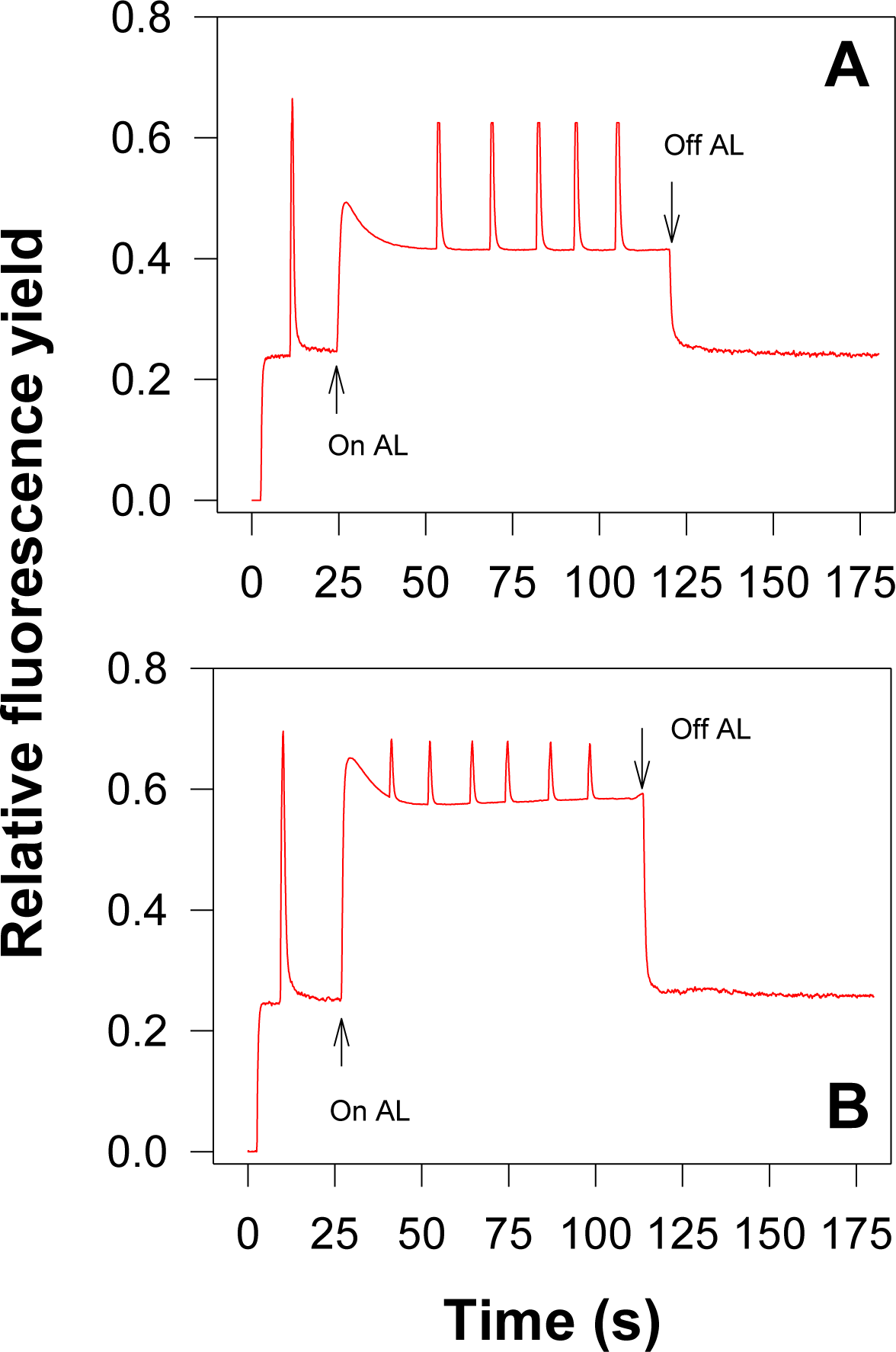
Chl fluorescence induction transients of the *Synechocystis* 6803 cell Δ*slr0201*strain) at room temperature. The dark-adapted cell suspensions were illuminated with saturating-pulse flashing to generate the maximal fluorescence yield (Fm), followed by actinic light (AL) switching-on to initiate photosynthesis, during which sequential saturating-pulse flashings were applied to monitor steady-state maximal fluorescence levels (Fm’).

State transitions in cyanobacteria reflect the redox state of the PQ pool. Several lines of evidence illustrated that PQ pool reductions induced a shift toward state 2 whereas its oxidations cause a state 1 transition (Mullineaux et al., 1989; Mao et al., 2002; Liu et al., 2012). Fig. 8 shows steady-state fluorescence kinetics (Fs, Fm and Fm’) of the wild type and the Δ*slr0201* mutant where, the dark-adapted cells were illuminated successively with either blue (Fig. 7A), or orange (Fig. 7B), or white light (Fig. 7C), which is in favor of PSI, PSII, and both PSII and PSI, respectively. The responses upon DBMIB treatments under different light illuminations were significantly different between the wild type and the Δ*slr0201* strain. Specifically, upon DBMIB addition under the blue-light illumination, the wild type exhibited a typical state 2 transition as Fm’ decreased consistently. The similar transition did not occur in the Δ*slr0201* strain. Under the orange-light illumination, the DBMIB-treated *slr0201* mutant cells demonstrated a shift from a transient, weak state 2 transition toward to a strong state 1 transition, reflecting a more oxidized PQ pool in the strain.

**Figure 7.**
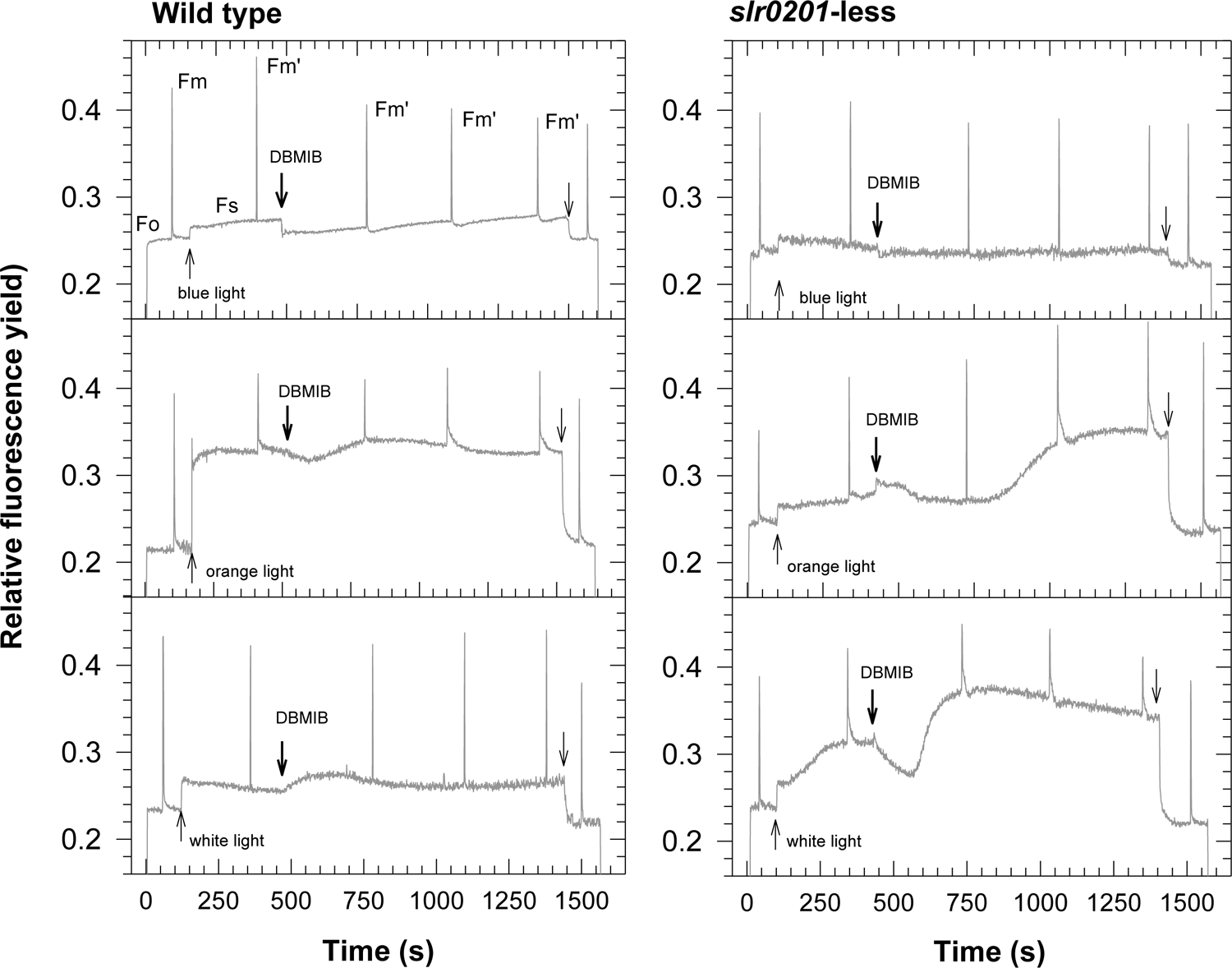
State transitions as fluorescence yield changes measured by a PAM fluorometer during different types of illumination in the wild-type and the Δstrain cells. The dark-adapted cells were first illuminated under non-actinic modulated light (M.L.) and followed by flashing using a saturating pulses 3500 μ s^−1^, 600 ms duration) to assess *F*_m_ dark (*F*_md_) and *F*_m_. After then, the cells were successively illuminated with either blue, or orange, or white light switching-On and switching-Off as shown, respectively by the up- and down thin arrows.

**Figure 8.**
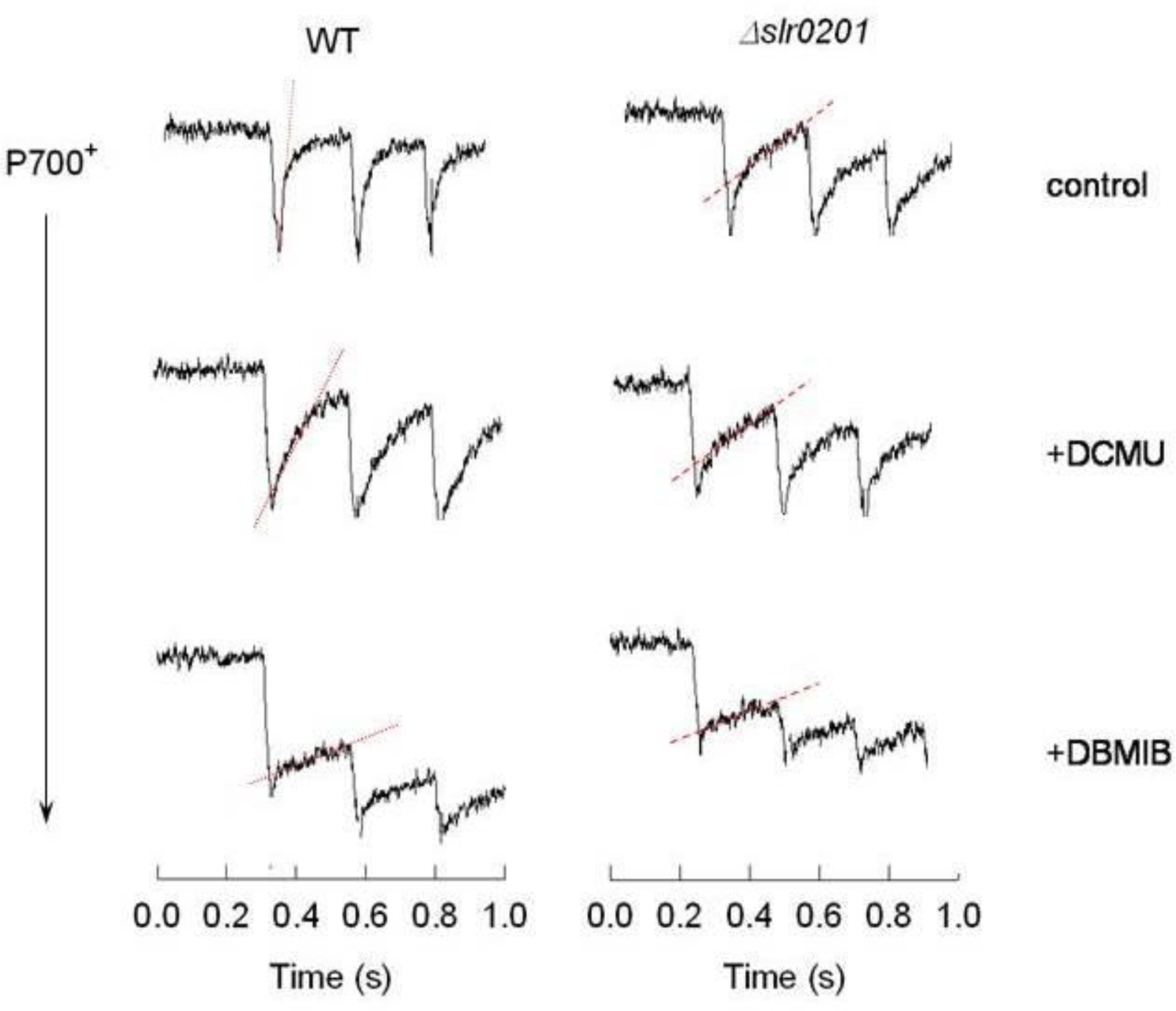
P700 oxidation-rereduction kinetics were affected by addition of electron transfer inhibitors that block either linear electron transfers from the PSII (DCMU) or respiratory electron transfer flows (DBMIB) through the PQ pool. Comparisons were made between the wild type and the *Δslr0201* strain using 100-ms flashings at a 250-ms interval to allow P700+ decay occurred. The slopes*, i.e.* the dotted and dashed lines, and the half-time were determined using an average from five flashing data.

### Post-illumination in vivo P700 re-reductions are delayed in the Δslr0201 strain

To further elucidate the redox state of the PQ pool as it may be affected by the impaired PSI-mediated cyclic electron transfer due to the Slr0201 knockout, kinetics of flash-on/off-induced P700 oxidation and rereduction were determined and compared between the wild type and the Δ*slr0201* mutant. Over a period of 10-300 ms, both P700 oxidation and P700^+^ rereduction increased with length of flashing illumination. The maximum amplitude reached at approximately 300 ms and 100 ms, respectively (data not shown). P700^+^ rereduction time-setting (100 ms) well fits in the range of recombination rate (∼ 90 ms ^-1^) between P700^+^ and F ^-^ in *Synechocystis* 6803 (Díaz-Quintana et al., 1998; Sétif et al., 2001). With a 100-ms flash illumination in absence of electron transfer inhibitors (Fig. 8), kinetics of flash-off induced Δsignals (P700 wild type was characterized by a dual-phase decay, i.e. a remarkably transient (∼ 25 ms) decay followed by a slow-phase decay over an extended period of 25-200 ms. The fast-phase decay with a slope of 9.5 ± 1.5 ms^-1^ (Fig. 8, the dotted line) accounting for ∼ 75% of overall rereduction of P700^+^ of which, the slow-phase decays contributed only ∼ 25%.

Compared to the wild type strain, the Δ*slr0201* showed not only a slower flash-induced photo-oxidation of P700 but also a markedly delay in P700^+^ rereduction with a half-time (t_1/2_) of 105 ms for the overall decay. Note that, although it was still evident, the transient fast-phase decay was almost diminished/ abolished such that it accounted for only ∼ 35% of the overall P700^+^rereduction. On the contrast, slow-phase decays were recorded, which accounted for a majority (∼ 65%) of the overall rereduction of P700^+^ with a slope of 0.6 ± 0.1 ms^-1^ (Fig. 8, the dashed line). As expected, addition of DCMU greatly delayed P700^+^ rereductions in both the wild type and the Δ*slr0201* strain due to abolishment of linear electron flows from the PSII. Under such circumstances, however, P700^+^ rereduction kinetics in the wild type was still much faster than that in the Δ*slr0201*.

Considering the fact that the slower reductions of P700^+^ in the Δ*slr0201* was attributed to the impaired respiratory electron flows into P700^+^ through the PQ pool. DBMIB treatment experiments seemed confirmed this notion. When electron flows coming out of the PQ pool were abolished by DBMIB, which will prevent P700^+^ rereductions via blocking both linear and cyclic electron transfers, only minim (leaking-mediated baseline) P700^+^ rereductions were monitored in both the wild type and the Δ*slr0201* mutant. Taken together, due to the impairments of PSI-mediated cyclic electron transfers, post-flashing P700^+^ rereductions in the Δ*slr0201* mutant occurred at much slower rates than in the wild type.

## Discussion

The primary aim of this study is the functional identification of the *slr0201*, an ORF in *Synechocystis sp.* PCC 6803 encoding a putative 301-amino acid protein that was originally annotated as HdrB. To achieve this goal, a *Δslr0201* mutant was constructed via knock-outing approach, followed by a series of physiological and biochemical phenotype analyses using the mutant. Based on the results from the study, it appears that the Slr0201 is a SDH-function-relevant protein rather than the HdrB as originally annotated (Kaneko et al., 1996). Several lines of evidence support this notion. Firstly, the original annotation of Srl0201 as HdrB was solely an output of sequence-based automatically-computational annotation without any phenotypic and/or functional analysis supports. In this context, while the Slr0201 does show high sequence similarities with HdrB, it also shows high sequence similarities with novel SdhCs from the ‘non-classical’ group of archaeal SDH that were recently found in archaeal species *Acidianus ambivalens* (Gomes et al., 1999; Lemos et al., 2001), *Sulfolobus acidocaldarius* (Janssen et al., 1997) and *S. tokodaii* (Iwasaki et al., 2002). In fact, HdrB also shares high sequence similarities with the novel Archaeal SdhC. For example, in *Acidithiobacillus ferrooxidans* ATCC 23270, an acidophilic, chemolithoautotrophic γ AFE_2550, a gene encoding HdrB was assigned SdhC, a succinate dehydrogenase C subunit (Quatrini et al., 2009). Secondly, as it is well known, functional HDR is a multiple-subunit complex (Hedderich et al., 1994; 2005). However, in the genome of *Synechocystis* 6803, except for a putative *hdr*B (*slr0201*) given that the original annotation were correct, no other *hdr*A/*hdr*C-like sequences are identified (Kaneko et al., 1996). This makes annotation of Slr0201 being HdrB largely speculative. Moreover, considering the fact that both *sdh*A-like (*slr1233*) and *sdh*B-like sequences (*sll0823*, *sll1625*) have been identified in *Synechocystis* genome, and functional involvements of these ORFs in succinate pathway were also experimentally confirmed (Cooley et al., 2000; Cooley and Vermaas, 2001), the absences of membrane anchoring subunit *sdh*C/D/E-like sequence in the genome of *Synechocystis* 6803 are obscure. Indeed, genome sequencing has indicated the SdhC-like protein in *Cyanobacterium aponinum* strain PCC10605 (Shiha et al., 2013)

In mitochondria or bacteria, SdhC usually serves as a membrane anchoring subunit associated with SDH complex (Lancaster, 2001; Iwasake et al., 2002; Schäfer et al., 2002). A sequence-based secondary structural analysis of the Slr0201was carried out and predicts the presence of at least one transmembrane helix in the hypothetic protein Slr0201, from 226 to 244, centered at 236 (data not shown), which is, in some cases, associated with a heme binding motif (*e.g.* James et al., 2005). A prokaryotic membrane lipoprotein lipid attachment site was also proposed for the residues 152-162. Whereas biochemical property of the Slr0201 awaits further identifications, the presence of transmembrane helix supports the notion that the Slr0201 acts as a membrane-anchoring subunit. This sequence-based structural prediction was confirmed by the overexpression experiments. The whole His-tagged Slr0201 overexpressed in *Synechocystis* 6803 was exclusively detected in the thylakoid membrane associated fragment (Xiong et al., 2014, unpublished).

Physiological analyses of the *Δslr0201* mutant as compared to the wild type and the Δ*sdhB_1_B_2_* mutant reveals some notably phenotypic characteristics. Probably the most interesting finding of the present study is that phenotypic characteristics of the Δ*slr0201* greatly resemble the Δ*sdhB_1_B_2_* mutant in terms of similar reduced respiratory rates (Table 1), impaired DCPIP reduction-based SDH activities (Fig. 4), lowed succinate and fumarate content (Table 1), reduced respiratory rates, and delayed KCN-induced fluorescence yield increases in the dark, an indication of a more oxidized PQ pool (Fig. 5, 6). Taken together, these results strongly indicate that the Slr0201 is a SDH function associated protein.

Among functional respiratory electron transport pathways in *Synechocystis* 6803, the SDH pathway contributes a large portion of electron flows into the PQ pool in the thylakoid membranes (Cooley et al., 2000; Cooley and Vermaas, 2001; Bernát et al., 2011). In the previous report, upon KCN addition in the dark, a quick, initial increase in fluorescence yield was monitored in the wild type cells, whereas in the Δmutant, only a markedly delayed, slow-phase rise in chlorophyll fluorescence was observed (Cooley and Vermaas, 2001). The result was interpreted as the impairments in the double mutant of the SDH-mediated cycle electron transfer around PSI. This observation was well refined in the KCN-treated Δ*slr0201* cells where, as Fig. 7 shows, the initial-phase increases in fluorescence yield were almost completely impaired. It is suspected that the delayed increases in fluorescence yield in the KCN-treated mutant cells might also be attributed to the impaired electron flux into the PQ pool through SDH-Δ*sdh*B-like phenotype favors the view that Δ*slr0201* encodes a SDH-relevant protein that is functionally involved in the interwoven ETCs in *Synechocystis* 6803.

Further evidence that Slr0201 is involved in SDH-mediated electron transfers around PSI was provided by fluorescence quenching and state transition analyses. The Δ*slr0201* mutant demonstrated remarkable impairments in both non-photochemical (qNP) and photochemical (qP) quenching due to the limited electron transfer capacities of the PSI. When compared to the wild type, the DBMIB-treated Δ*slr0201*mutant exhibited a relatively weak state 2 transition under blue light and a strong state 1 transition under orange light (Fig. 7). These results are favorable to the view that *slr0201* knock-outing impaired SDH-mediated cycle electron flows around PSI and resulted in a more oxidized PQ pool in the mutant. P700^+^ rereduction kinetics provides further insight into cyclic electron transfers around PSI, which was balanced by SDH pathway-mediated electron flows through the PQ pool. With addition of DCMU to prevent the PSII-driven electrons from reaching the Cyt-*b_6_*f complex and thereby masking any rate limiting effect on the acceptor side of PSI, electrons flow available for P700^+^ rereduction will be then predominantly derived from NADH dehydrogenase (Mi et al., 1992) and/or SDH (Cooley et al., 2000; Cooley and Vermaas, 2001; Bernát et al., 2011). The Δ*slr0201* mutant showed not only a slower flash-induced photo-oxidation of P700 but also a delayed rereduction with a t_1/2_ of 105 ms^-1^ significantly slower than in the wild type (Fig. 8).

HPLC-GC/MS analysis of the organic acid contents in the *slr0201*mutant provides evidence for the SDH-pathway associated Slr0201. Fumarate level decreased significantly in the mutant, which is consistent with the expected consequence of an impaired SDH function. However, the decreased fumarate levels in the mutant cells were not essentially associated with increases in succinate level (Table 1; see also Cooley and Vermaas, 2001). This indicates that the cells may possess certain kinds of unknown mechanisms to adjust succinate level accordingly. For example, it would be possible that when the conversion from succinate to fumarate was down-regulated, other succinate utilization pathways, such as glutamate metabolism, may be activated and up-regulated in order to feedback biosynthesis of related metabolites. The availability of succinate might be a rate-limiting factor in respiratory electron transfer considering the fact that SDH-mediated respiratory ET shows larger electron donation capacity for reduction of the PQ pool (Cooley et al., 2000).

L-arginine catabolism in cyanobacteria involves cyanophycin synthesis which is regulated by not only nutrient deficiency but also photosynthetic activity, especially the activity status of PSII. A PsbO-free mutant with reduced excitation pressure on PSII due to lack of manganese stabilizing peptide were able to maintain balanced grow when cultivated on arginine as sole N-source (Stephan et al., 2000). Compared to the wild type whose growths were largely reduced with presence of L-arginine as sole N-source, the PsbO-free mutant showed substantial impairments in the PSII activity. Since a reduced excitation pressure on PSII favors L-arginine catabolism to maintain a balanced growth and the impairments in growth occurred when PSI activity was impaired, it is assumed that growth sensitivities of the Δ*slr0201* mutant to arginine might be attributed to the impaired PSI activity due to the reduced SDH-mediated cyclic electron transfer. In the Δ*slr0201* mutant, impairments in the PSI function may result in, as the plate and liquid culture results indicated (Figs. 2 and 3) generation and accumulation of certain kinds of toxic chemical and /or metabolites, which warrants further experimentation to identify and elucidate the details.

In summary, the *slr0201* knockout strain was physiologically characterized by a *ΔsdhB_1_B_2_*-like phenotype with pronounced impairments of the SDH-relevant functions that include reduced SDH activity, impaired dark respiration, lower succinate and fumarate content, more oxidized PQ pool in darkness, and delayed KCN-induced transient increases of fluorescence yield in darkness. While these results strongly support the hypothesis that SDH-mediated respiratory electron flows functionally involve in the redox poising of PQ pool in thylakoid membrane in cyanobacterium, physiological characterizations of the *Δslr0201* in combination with sequence aligning prediction suggest that the ORF *slr0201* encodes a SDH-function-relevant protein, rather than heterodisulfide reductase B subunit (HdrB). The putative Slr0201 acts as a membrane-anchoring subunit of SDH complex in *Synechocystis* 6803. Sequence analysis indicates that the Slr0201 is highly homologue to the SdhC in the ‘non-classical’ group of SDH that were found in certain archaeal species (Lemos et al., 2001; Iwasaki et al., 2002; Schäfer et al., 2002). This implies that the *Synechocystis* 6803 SDH may belong to Type V of succinate dehydrogenase which is genetically distinct from mitochondrial or bacterial SdhC (Lancaster, 2001; Lemos et al., 2002).

## Acknowledgements

I would like to thank Dr. Karl Booksh for access to the GC/MS equipment. The funding support for this research was provided by a grant to WV from the Department of Energy (DE-FG03-01ER15251).

## Notes

### Competing Interest Statement

The authors have declared no competing interest.

